# First report of *Theileria annulata* in Nigeria: findings from cattle ticks in Zamfara and Sokoto States

**DOI:** 10.1101/2020.11.10.376624

**Authors:** Adamu Haruna Mamman, Vincenzo Lorusso, Babagana Mohammed Adam, Abraham Goni Dogo, Kevin J Bown, Richard J Birtles

## Abstract

**Background:** Ticks and tick-borne pathogens (TBPs) represent a significant economic burden to cattle farming in sub-Saharan Africa including Nigeria. However, in the northern part of this country, where the largest livestock population reside, little is known about the contemporary diversity of ticks and TBPs. This area is particularly vulnerable to climate change, undergoing marked transformation of habitat and associated flora and fauna that is also likely to include ticks. This study aimed to document the occurrence of tick species and Apicomplexan TBPs in cattle from North-Western Nigeria.

**Methods:** In 2017, ticks were collected from cattle in Zamfara and Sokoto States and identified morphologically. Additionally, a subset of ticks were screened molecularly for the detection of Apicomplexan DNA.

**Results:** A total of 494 adult ticks were collected from 80 cattle in Zamfara and 65 cattle in Sokoto State. Nine tick species were encountered, including seven *Hyalomma* spp. (i.e. *Hyalomma dromedarii, Hyalomma impeltatum, Hyalomma impressum, Hyalomma marginatum, Hyalomma rufipes, Hyalomma truncatum* and *Hyalomma turanicum*), *Amblyomma variegatum* and *Rhipicephalus* (*Boophilus*) *decoloratus*. All species were present in Zamfara, whereas only five species were found in Sokoto. *Hyalomma rufipes* was the most prevalent tick infesting cattle in Zamfara State (76.2%), while *H. dromedarii* was the most prevalent in Sokoto State (43.7%), confirming the widespread transfer of this species from camels onto livestock and its adaptation to cattle in the region.

Of 159 ticks screened, 2 out of 54 (3.7%) from Zamfara State and 29 out of 105 (27.6%) from Sokoto State harboured DNA of *Theileria annulata*, the agent of tropical theileriosis.

**Conclusions:** This study confirms the presence of a broad diversity of tick species in cattle from North-Western Nigeria, providing the first locality records for Zamfara State. The occurrence of *H. turanicum*, recorded for the first time in Nigeria, indicates a distribution of this tick beyond Northern Africa.

This study provides the first report for *T. annulata* in Nigeria. Given its enormous burden on livestock farming in North Africa and across Asia, further investigations are needed to better understand its epidemiology, vector transmission and potential clinical significance in cattle from Northern Nigeria and neighbouring Sahelian countries.

## Background

Ticks represent a significant economic burden to cattle farming and, overall, the development of the livestock sector in sub-Saharan Africa (SSA) [1, 2, 3]. Their significance is due to the impairment they cause to livestock productivity, attributable to both the direct and indirect effects of tick’s parasitism and blood feeding [2]. In cattle, direct damage caused by ticks include anaemia, stress (‘tick worry’), reduction of feeding thus decrease of weight gain and milk yields, susceptibility to secondary infections, devaluation of hide quality, hypersensitivity and toxicosis [1, 2]. Indirect adverse consequences of tick infestation in cattle are linked to the conditions that are caused by the plethora of tick-borne pathogens (TBPs), including mostly protozoa and bacteria, but also helminths, viruses and fungi, some of which are of zoonotic importance [1–4]. The most important TBPs threatening cattle health and productivity in SSA are the causative agents of theileriosis (i.e. the ‘East Coast fever’ agent *Theileria parva; Theileria annulata; Theileria mutans* and *Theileria velifera*), babesiosis (i.e. *Babesia bigemina* and *Babesia bovis*), anaplasmosis (i.e. *Anaplasma marginale, Anaplasma centrale* and *Anaplasma bovis*) and ehrlichiosis (i.e. the ‘heartwater’ agent *Ehrlichia ruminantium*) [1].

In spite of the enormous burden of ticks and TBPs on livestock farming, for many parts of SSA, even fundamental epidemiological information is lacking. Nigeria is a case in point; despite one of the largest cattle populations in the continent (of approximately 20 million heads) [5], contributing one third of national agricultural GDP and providing 36.5% of the total protein intake of Nigerians [6], substantial gaps affect the current understanding of the epidemiology of ticks and TBPs in the country [7], with knowledge of cattle-associated tick diversity and distribution being rather patchy [8–12] when not outdated [13–16]. Additionally, although approximately 90% of the country’s cattle population is concentrated in the Northern region [6, 17], most historical surveys were carried out in southern States [13, 18]. So far, published investigations on cattle ticks from Northern Nigeria have focused on limited areas of eastern (e.g. Maiduguri and Yobe State) [10–11], or western States (e.g. Sokoto and Kebbi States) [19–22], limiting in some instances the identification of ticks to the genus level [10, 11, 20]. Similarly, the majority of studies on TBPs in Nigeria, detecting the presence of apicomplexan parasites belonging to the genera *Theileria* and *Babesia*, and members of the bacterial genera *Anaplasma, Ehrlichia, Rickettsia* and *Coxiella* spp., have mostly relied on cytological (i.e. microscopical examination of blood smears and biopsies) [23–28] and serological approaches (e.g. ELISA and immunofluorescence assays) [26, 29–32], and only in a few, recent instances on the molecular screening of bovine blood [33–35] and ticks [36, 37]. Moreover, to date no study based on molecular diagnostic techniques has ever been carried out in the North-Western region.

The epidemiological importance of surveying ticks and TBPs in cattle from Northern Nigeria is also enhanced by the frequent movement and introduction in this region of livestock hailing from neighbouring countries like Niger, Chad and Cameroon, brought to Nigeria to be sold in more profitable local markets [38]. Furthermore, the heavy reliance on climate-sensitive economic activities, such as agriculture and livestock keeping, makes Northern Nigeria particularly vulnerable to climate change [39]. Spanning the Sudano-Sahelian ecological zone [40, 41], this region is currently experiencing a combination of rising heat and declining rainfall that together are accelerating desert encroachment and marked habitat change [39, 42, 43]. Besides affecting cattle health directly through their effects on water and pasture availability, these alterations may also lead to indirect negative consequences, linked to the likely changes that they will cause on tick populations’ diversity and ecology [44]. Habitat changes may indeed compromise the fitness of some endemic tick species and create new niches exploitable by exotic species, originating from Sahelian and North African countries, adapted to hot and dry environments. The arrival of such species may well be accompanied by the TBPs they vector.

The present study aimed therefore to determine the contemporary diversity of cattle-associated ticks and apicomplexan TBPs in a region of North-Western Nigeria heavily reliant on cattle keeping and significantly affected by climate change [43, 45, 46], with the objective of assessing the extent of change that maybe be attributed to the latter’s impact.

## Methods

### Study area

Field activities were carried out between March and May 2017, in two north-western States of Nigeria, namely Zamfara and Sokoto, where a convenience sample of cattle were inspected and surveyed for tick infestations. In Zamfara, ticks were collected from cattle in the villages of Anka, Kwaye, Kwakwalwa, Gema and Abara, all of which lie within the Anka Local Government Area (LGA) (11°59’N and 6°02’E). In Sokoto, ticks were collected from cattle in cattle markets from three LGAs, namely Sokoto North, Wurno, and Illela (13° 03’N and 5° 14’E) (Table 1).

**Table 1.** Number of animals surveyed, number of ticks collected and mean tick loads.

| State | Local Government Area (LGA) | Village | No. of Cattle sampled | Ticks collected | | | Mean tick count/animal ± SE | |
| --- | --- | --- | --- | --- | --- | --- | --- | --- |
|  |  |  |  | Males | Females | TOTAL | Village level | State level |
| Zamfara | Anka | Anka | 16 | 36 | 6 | 42 | 2.62 ± 1.67 | 3.0 ± 2.28* |
| | | Kwaye | 14 | 22 | 14 | 36 | 2.57 ± 2.31 | |
| | | Kwakwalwa | 11 | 28 | 15 | 43 | 3.9 ± 1.87 | |
| | | Gema | 15 | 32 | 14 | 46 | 3.06 ± 2.12 | |
| | | Abara | 24 | 56 | 17 | 73 | 3.04 ± 2.88 | |
| Sub-total |  |  | 80 | 174 | 66 | 240 |  |  |
| Sokoto | Sokoto North | Kara | 13 | 18 | 36 | 54 | 4.15 ± 4.20 | 3.91 ± 3.15* |
| | Wurno | Achida | 20 | 40 | 54 | 94 | 4.70 ± 3.87 | |
| | Illela | Illela | 32 | 29 | 77 | 106 | 3.31 ± 1.91 | |
| Sub-total |  |  | 65 | 87 | 167 | 254 |  |  |
| TOTAL |  |  | 145 | 261 | 233 | 494 |  |  |

Both Sokoto and Zamfara are among the poorest States of the Federal Republic of Nigeria [47]. Their economy is almost entirely reliant on agriculture. Livestock including cattle, sheep and goats are reared, and some crops are grown. Donkeys and camels are commonly used as draft animals [45, 48].

The region lies on the boundary of Sudan savanna and Sahel climatic zones [41]. The meteorology is seasonal, with a 3-4 month wet season occurring between June and September, during which time about 500 mm of rain falls. The remainder of the year is very dry. Average daily temperatures range between about 18 °C and 38 °C. The vegetation is characterised as open savanna grasslands or open savanna woodland, with fine-leaved and broad-leaved trees and shrubs, which are deciduous for several weeks [49].

### Tick collection and identification

Each animal sampled was restrained by its owner/herder and its hide examined carefully, focusing in particular on established predilection sites for tick attachment (i.e. ear, dewlap, abdomen, hooves, inguen, perineum, peri-anal region, tail) [8, 50]. From these anatomical regions, all visible adult ticks were collected using steel forceps to remove each specimen in its entirety. Immediately after collection, all ticks removed from the same individual cattle were placed in a 5 ml plastic tube containing 70% ethanol, before being transported to the University of Salford for further analysis. Once in the laboratory, all collected ticks were identified to the species level on the basis of observed anatomical features, using the taxonomical keys by Walker et al. [51].

### Detection and identification of Apicomplexa using molecular methods

A subset of collected ticks (n=159, 32.2% of total ticks) were screened for apicomplexan pathogens using molecular methods. These ticks were chosen to embrace all species encountered and the different locations in which each tick species was encountered (Table 3) Crude DNA extracts, prepared from individual ticks as previously described [52], were incorporated into a previously described PCR targeting an 18S rDNA fragment specific to apicomplexan taxa [53]. PCRs were prepared in a dedicated DNA-free laboratory. “Blanks” (PCRs containing water instead of DNA extracts) were co-processed with all samples at a ratio of 5 samples:1 blank, to test for cross-contamination. Reagent controls (a DNA-free negative and a *Babesia microti* positive) were also included in each set PCRs prepared. The success of the PCR was assessed by UV visualisation of GelRed-stained amplification products (of about 680 base pairs) following their electrophoretic resolution on a 1% (w/v) agarose gel. Amplification products were purified using an Isolate II PCR and gel kit according to manufacturer’s instructions (Invitrogen, Carslbad, California, USA). Sanger sequencing of both strands of each PCR product was carried out commercially. Chromatograms obtained were visualised using Chromas Pro software (Technelysium, Brisbane, Australia). Data from complementary strands of each amplicon were aligned with one another and regions of ambiguity together with primer sequences at the extremities were removed. The identity of the organism from which a sequence obtained was determined by comparison with data held on GenBank using Basic Local Alignment Search Tool (BLAST).

### Statistical analysis

Data were entered in Microsoft Excel, through which mean tick infestations and standard errors were calculated at the study village, LGA and State (i.e. Zamfara and Sokoto State) level (Table 1). For the two States, mean prevalence, including 95% Confidence Intervals (CI), of tick species retrieved were calculated using the WinPepi software (Version 11.6). Using the same software, cumulative tick counts recorded for each State were compared statistically using the Chi-squared test. In addition, infection rates were compared according to tick species and State of provenance (for both ticks and cattle), using the two-tailed Fisher’s exact test. P values <0.05 were considered statistically significant.

## Results

### Tick identification and infestation burden

In total, 494 adult ticks were collected from 145 cattle; of these 254 off 65 cattle in Sokoto and 240 off 80 cattle in Zamfara (Table 1). The mean infestation rate was 3.0 in Zamfara and 3.9 in Sokoto, with no statistically significant difference being recorded (P=0.8) (Table 1). A total of nine tick species were encountered, these included seven *Hyalomma* species (i.e. *Hyalomma dromedarii, Hyalomma impeltatum, Hyalomma impressum, Hyalomma marginatum, Hyalomma rufipes, Hyalomma truncatum* and *Hyalomma turanicum*), *Amblyomma variegatum* and *Rhipicephalus* (*Boophilus*) *decoloratus* (Figure 1, Table 2). All nine species were present in Zamfara and these included *H. turanicum* (Figure 1.VII, a-b), recorded for the first time in Nigeria. Only five species were present in Sokoto, namely (from the most to the least prevalent) *H. dromedarii*, *H. impeltatum*, *H. truncatum, H. impressum* and *H. rufipes* (Table 2). However, these corresponded to five of the six most abundant species collected in Zamfara, with only those species for which only a single specimen was found (i.e. *H. marginatum, H. turanicum* and *Rh. (Bo.) decoloratus*) and *A. variegatum* being absent (Table 2).

**Figure 1.**
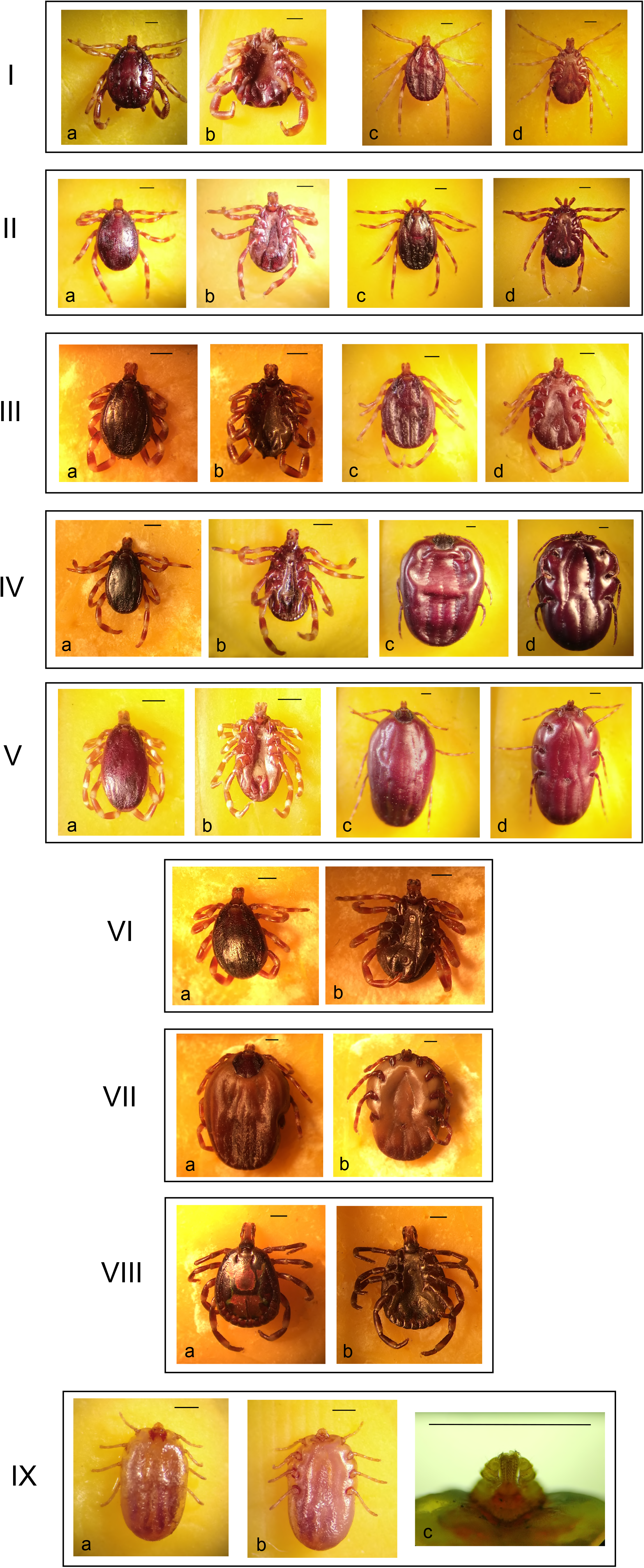
Tick species encountered in this survey. From top to bottom: *Hyalomma dromedarii* (**I**): adult male, dorsal and ventral view (**I, a-b**), and adult female, dorsal and ventral view (**I, c-d**); *Hyalomma rufipes* (**II**): adult male, dorsal and ventral view (**II, a-b**), and adult female, dorsal and ventral view (**II, c-d**); *Hyalomma impeltatum* (**III**): adult male, dorsal and ventral view (**III, a-b**), and adult female, dorsal and ventral view (**III, c-d**); *Hyalomma truncatum* (**IV**): adult male, dorsal and ventral view (**IV, a-b**), and adult female, dorsal and ventral view (**IV, c-d**); *Hyalomma impressum* (**V**): adult male, dorsal and ventral view (**V, a-b**), and adult female, dorsal and ventral view (**V, c-d**); *Hyalomma marginatum* (**VI**): adult male, dorsal and ventral view (**VI, a-b**); *Hyalomma turanicum* (**VII**): adult female, dorsal and ventral view (**VII, a-b**); *Amblyomma variegatum* (**VIII**): adult male, dorsal and ventral view (**VIII, a-b**); *Rhipicephalus (Boophilus) decoloratus* (**IX**): adult female, dorsal and ventral view (**IX, a-b**), and details of the ventral view of the mouthparts (**IX, c**) showcasing 3 + 3 rows of hypostomal teeth and the protuberance with pectinate setae on the internal margin of palp article I. Black bar = 1 mm.

**Table 2.** Cumulative counts, prevalence, number of males and females, and male: female ratio of ticks identified.

| State | Tick species | Total | Mean Prevalence %<br>(95% confidence interval) | Males | Females | Male : Female ratio |
| --- | --- | --- | --- | --- | --- | --- |
| Zamfara | *Hyalomma rufipes* | 183 | 76.2 (70.3 - 81.5) | 135 | 48 | 2.8 : 1 |
|  | *Hyalomma truncatum* | 18 | 7.5 (4.5 - 11.6) | 16 | 2 | 8 : 1 |
|  | *Hyalomma impeltatum* | 14 | 5.8 (3.2 - 9.6) | 6 | 8 | 1 : 1.3 |
|  | *Hyalomma impressum* | 9 | 3.7 (1.7 - 7.0) | 8 | 1 | 8 : 1 |
|  | *Amblyomma variegatum* | 7 | 2.9 (1.2 - 5.9) | 7 | 0 | 7 : 0 |
|  | *Hyalomma dromedarii* | 6 | 2.5 (0.9 - 5.3) | 1 | 5 | 1 : 5 |
|  | *Hyalomma marginatum* | 1 | 0.4 (0.0 - 2.3) | 1 | 0 | 1 : 0 |
|  | *Hyalomma turanicum* | 1 | 0.4 (0.0 - 2.3) | 0 | 1 | 0 : 1 |
|  | *Rhipicephalus (Boophilus) decoloratus* | 1 | 0.4 (0.0 - 2.3) | 0 | 1 | 0 : 1 |
| **Total** |  | **240** |  | **174** | **66** | **2.6 : 1** |
| Sokoto | *Hyalomma dromedarii* | 111 | 43.7 (37.5 - 50.0) | 47 | 64 | 1 : 1.4 |
|  | *Hyalomma impeltatum* | 93 | 36.6 (30.7 - 42.9) | 29 | 64 | 1 : 2.2 |
|  | *Hyalomma truncatum* | 27 | 10.6 (7.1 - 15.1) | 7 | 20 | 1 : 2.9 |
|  | *Hyalomma impressum* | 15 | 5.9 (3.3 - 9.5) | 0 | 15 | 0 : 15 |
|  | *Hyalomma rufipes* | 8 | 3.1 (1.4 - 6.1) | 4 | 4 | 1 : 1 |
| **Total** |  | **254** |  | **87** | **167** | **1 : 1.9** |

There was a marked difference between the relative abundance of tick species in each State. In Zamfara, *H. rufipes* dominated (76.2%), with no other species accounting for more than 8% of samples. Conversely, in Sokoto, *H. dromedarii* was most abundant (43.7%) and, together with *H. impeltatum*, accounted for 80% of the ticks collected (Table 2). *Hyalomma rufipes* was present in Sokoto, but at a relative prevalence of only 3.1%, whereas *H. dromedarii* was present in Zamfara, but only at a relative prevalence of 2.5% (Table 2). Overall 261 male ticks and 233 female ticks were collected. The ratio of male to female ticks was very different in the two States, being 2.6:1 in Zamfara and 1:1.9 on Sokoto (Table 2).

### Molecular screening

Out of the total of 159 ticks screened molecularly, 31 ticks (19.5%) were found positive for Apicomplexan DNA, with all bar two being ticks from Sokoto (Table 3). Significantly more ticks from Sokoto (29/105 tested, 27.6%) yielded a PCR product than ticks from Zamfara (2/54, 3.7%) (P<0.001) and significantly more cattle from Sokoto (18/35 tested, 51.4%) bore infected ticks than cattle from Zamfara (2/35 tested, 5.7%) (P<0.001).

**Table 3.**
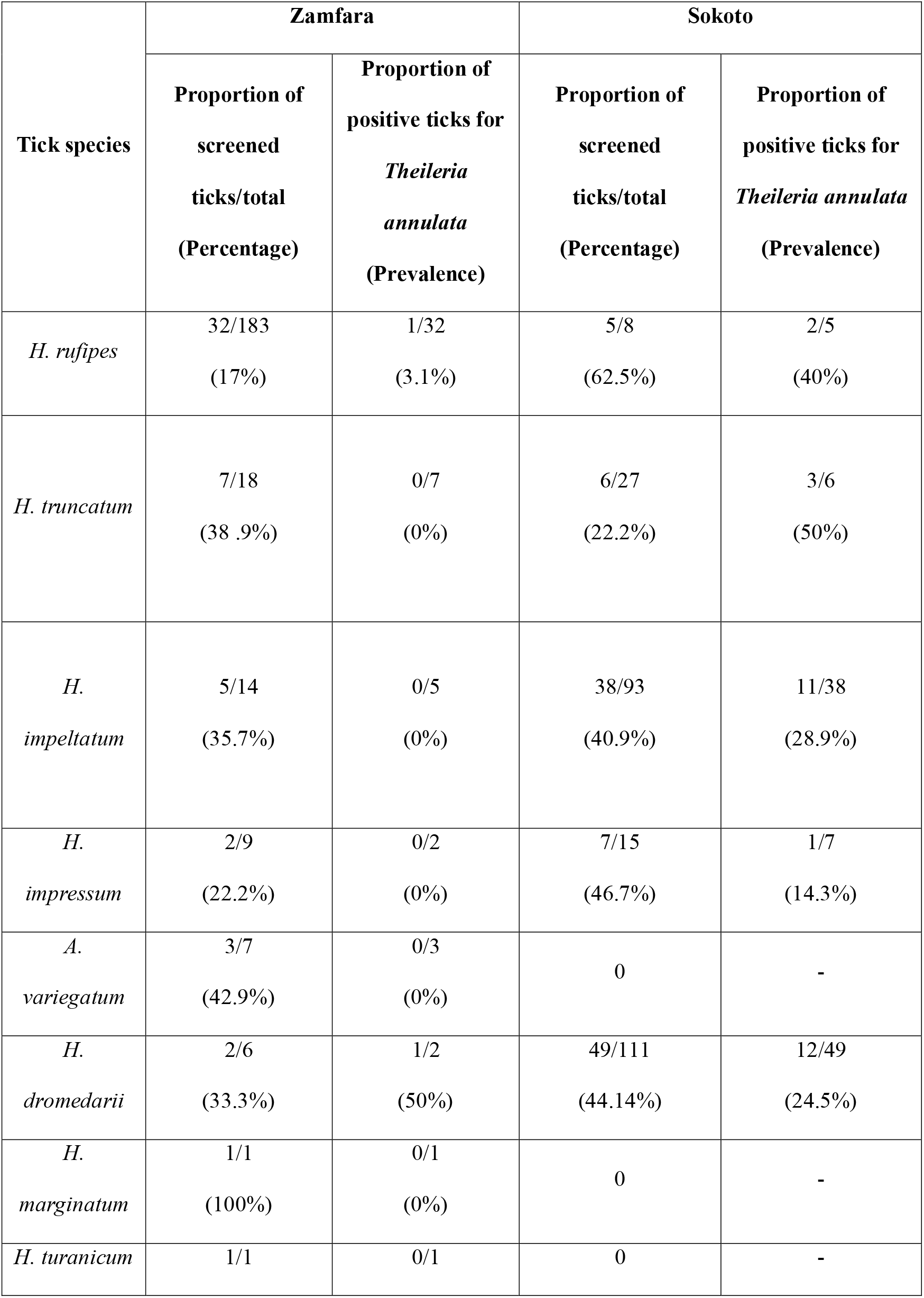
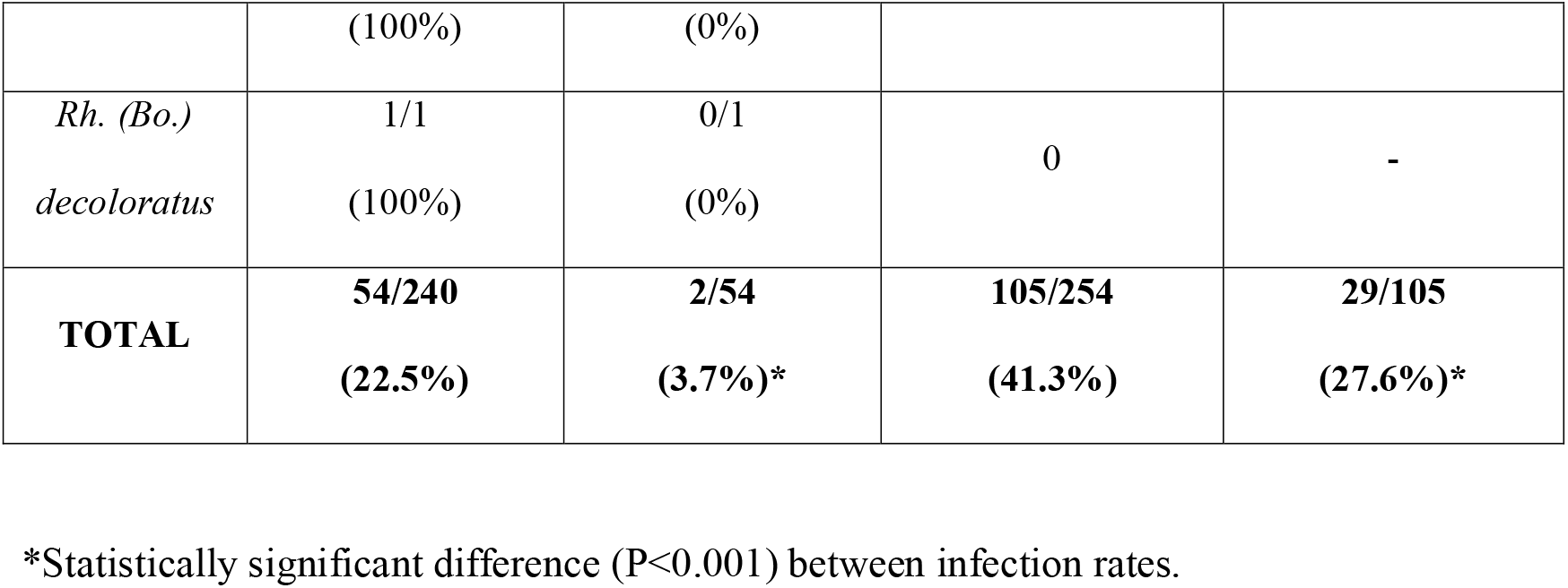
Ticks screened and tested positive for the detection of Apicomplexan (i.e. *Theileria annulata*) DNA.

Unambiguous sequence data were obtained from 23 of the amplicons, including those obtained from the two positive ticks from Zamfara. BLAST analysis of these sequences (609 base pairs) revealed all to be indistinguishable from one another and to share 100% similarity with partial 18S rDNA sequences of many strains of *Theileria annulata* (e.g. GenBank MN944852). The 18S rDNA sequence obtained shared less than 99.4% similarity with those from other *Theileria* species (most similar was 4 SNPs compared to *Theileria lestoquardi* 18S rDNA, GenBank AF081135). A selected sequence amongst those obtained was deposited in GenBank on 30 October 2020 (GenBank MW191850).

Tick species containing *T. annulata* DNA included mainly *H. dromedarii* and *H. impeltatum*, but also *H. truncatum, H. impressum* and *H. rufipes* (i.e. all species encountered in Sokoto) (Table 3).

The prevalence of infection in each tick species present in Sokoto did not vary significantly (P=0.6), ranging from 14.3% in *H. impressum* to 50% in *H. truncatum* (Table 3). The large majority (n=27/31; 87.1%) of PCR positive ticks were females, assessed as being either partially (n=16) or very engorged (n=11) (data not shown). However, four PCR positive ticks were male, one of which (i.e. *H. dromedarii*) appeared to be unfed.

## Discussion

All cattle surveyed in this study were infested by adult ticks, in both Zamfara and Sokoto States. The generally relatively low degree of infestation, which did not differ significantly between the two States, could be the outcome of (i) the time of the year when the sampling took place (i.e. late dry season, when most tick populations are expected to be less abundant than in the wet season) and (ii) the traditional control practice based on manual removal of ticks, adopted periodically (e.g. once a week) by cattle keepers [8].

The rich diversity of tick species parasitising cattle encountered in this survey is generally in keeping with contemporary reports in Nigeria [8, 9]. As expected for sites in the Sahel, *Hyalomma* species dominated [51, 54–57]. While it represents the first investigation of this kind for the State of Zamfara, this study confirms the occurrence of five *Hyalomma* spp. (i.e. *H. dromedarii*, *H. impeltatum*, *H. impressum* and *H. trunctatum*) which were previously recorded, although in different prevalences, by two surveys carried out on cattle in Sokoto [19, 21].

The presence of such a high relative prevalence of *H. dromedarii* is remarkable. Commonly known as ‘the camel *Hyalomma*’, this tick has a wide distribution in Africa, encompassing arid and desertic regions that are north of the equator, where it is the predominant species parasitising camels [51]. Numerous previous reports demonstrate its ability to infest several hosts including cattle, but not with the relative success observed in this study [54, 58]. In Sokoto, *H. dromedarii* was the most frequently encountered tick, accounting for almost half the total number of collected ticks (43.7%), suggesting frequent transfer of ticks from camels to cattle and/or that *H. dromedarii* has adapted itself to parasitise cattle here to a degree not reported elsewhere in its range. The presence of *H. dromedarii* on cattle in the absence of (frequent) camel contact has indeed been noted previously but only at very low infestation rates [59]. Our findings are in general agreement with those from two previous surveys carried out on cattle from Sokoto, in 2009-10 [19] and 2013 [21], recording *H. dromedarii* as the second the most prevalent tick species (13.3% and 15.4%, respectively), in both cases after *H. truncatum* (15.5% and 18.4%, respectively) [19, 21]. Nonetheless, the remarkably higher prevalence documented by the present investigation may be suggestive of adaptation of *H. dromedarii* towards parasitising cattle which may have occurred in recent times in Sokoto State. Further investigations aiming to verify this hypothesis and assess the potential implication of ecological and climatic factors, would be advisable.

Our observation of the dominance of *H. dromedarii* on cattle in Sokoto mirrors the findings by Lawal et al. [56] and Onyiche et al. [57], reporting *H. dromedarii* as the most prevalent (46.9%) [56] and the second most prevalent (42.3%) [57] tick species collected in camels from Sokoto. Interestingly, in the very recent survey by Onyiche et al. [57], specimens of *H. dromedarii* tested positive for the presence of DNA of the Q fever agent *Coxiella burnetii*. The fact that the dominance of *H. dromedarii* on cattle was not also observed in Zamfara suggests that the ecological niche it currently occupies in Sokoto does not extend southward within the Sudan savanna of Nigeria and could also be attributable to Zamfara’s smaller camel population compared to Sokoto [59].

In Zamfara, *H. rufipes* was by far the most abundant tick encountered. This species is the most widespread member of the genus present in Africa and its major contribution to cattle-associated tick fauna has been reported at several sites across this range [55, 60, 61]. It was previously identified in cattle from North-Central (i.e. Plateau State) [8, 9], North-Eastern (i.e. Taraba and Borno States) and North-Western Nigeria (i.e. Kaduna, Kano, Katsina and Sokoto States) [9, 19, 21]. Its veterinary importance is linked to its capacity to act as vector of *Anaplasma marginale* [62] and *Babesia occultans* [63]. It is also recognised as an important vector Crimean-Congo haemorrhagic fever (CCHF) virus to humans [51]. Importantly, specimens of *H. rufipes* collected from camels at slaughterhouses in Kano State, were previously found positive for DNA of *Rickettsia aeschlimannii*, responsible for a zoonotic spotted fever [66].

*Hyalomma impeltatum* was the second most prevalent tick species (n=93/254; 36.6%) in Sokoto State; this tick was found in more modest numbers in Zamfara State (n=14/240; 5.8%). Commonly infesting camels in Northern Nigeria [56, 64], this species was also documented in trade cattle that reached Ibadan, South-Western Nigeria, hailing from the North of the country or neighbouring Chad or Niger [27] as well as in cattle from Sokoto State [19, 21]. The prevalence of *H. impeltatum* recorded by this survey in Sokoto (i.e. 36.4%) was higher than that previously documented in cattle from the same State (10.1% and 9.4%, respectively) [19, 21], confirming the adaptability of this species to parasitising cattle in areas of sympatry with camels. Like for *H. rufipes*, specimens of *H. impeltatum* collected from camels in Kano State were found to harbour DNA of the zoonotic *R*. *aeschlimannii* [64]. In this study, specimens of *H. truncatum* were recorded in cattle from both Zamfara and Sokoto State (Table 2). This tick is known for commonly infesting cattle in Nigeria [8, 9, 15, 16, 19, 21]; its significance is mostly related to its capacity to cause a toxic syndrome (‘sweating sickness’), particularly in young cattle [65] and to the injuries caused by its long mouthparts, especially in the interdigital clefts [51]. Like *H. dromedarii, H. truncatum* specimens collected from camels from Northern Nigeria (i.e. Sokoto, Kano and Jigawa States) were found to harbour *C. burnetii* DNA [57].

This study confirms the occurrence of *H. impressum* in North-Western Nigeria. Initially documented in trade in Ibadan [27], this tick was previously described in cattle from Sokoto State [19, 21]. The fact that *H. impressum* was encountered on cattle in this study but not in the two aforementioned surveys on camels from Sokoto [56, 57] may reflect a stronger host-specificity for cattle, although this conclusion is confounded by the low relative abundance of this species on the cattle we observed. Moreover, specimens identified as *H. impressum* were also encountered in camels in Kano State [56, 64]. Very little is known about the veterinary importance and vectorial competence of this tick species, although some engorged specimens in Ibadan showed to harbour kinetes of ‘*Babesia* spp.’ [27].

One male specimen of *H. marginatum* was identified in cattle from Zamfara State. It was morphologically distinguished from the closely related *H. rufipes*, especially on the basis of the distinctive pattern of grooves on its conscutum. Usually reported in livestock across the Maghreb region, Egypt, Sudan and Ethiopia [51], the occurrence of this tick species may be more common that known until now in the Sahel region. In Nigeria, this tick was previously detected in trade cattle probably introduced from neighbouring Chad or Niger [27]. It is however thought of being unable to survive under desert conditions [50], which may explain its absence from cattle surveyed in Sokoto State, in this study as well as in previous ones [19, 21] and suggest that it may be more prevalent during the wet season months. Like *H. rufipes*, it can be a vector of the zoonotic CCHF virus [51].

Perhaps the most noteworthy encounter among cattle-associated ticks in Zamfara was *H. turanicum*. This species has, to our knowledge, not been reported in Nigeria, or elsewhere in West Africa previously. It is thought to be endemic in the north-east of the continent, and is established in arid, hot parts of southern Africa after accidental introduction [51]. This tick is not known to transmit pathogens to livestock, although it is considered a vector of the CCHF virus to humans [51]. *Hyalomma turanicum* has a two-host life cycle, with adults typically parasitising wild and domesticated large ruminants and larvae and nymphs feeding on smaller mammals and ground-frequenting birds [51]. The tick has also been reported in Europe, associated with migratory birds using the western European-African flyway [66]. As this flyway embraces Nigeria and large parts of Africa north of the Sahara, it is reasonable to propose that the *H. turanicum* observed in Zamfara was introduced as a feeding nymph by a migratory bird. As yet to it too early to conjecture if *H. turanicum* is established in Northern Nigeria; further surveys of cattle and likely hosts of immature life-stages would help clarify this uncertainty. Considering the suitability of this tick species for desert and steppe landscapes [51], climate change-induced desert encroachment in the Sahelian region may potentially favour the establishment of *H. turanicum* in Northern Nigeria.

A few (n=7), all male specimens of *A. variegatum* were recorded in cattle from Zamfara. The low numbers of *A. variegatum* recorded in Zamfara, coupled with its absence from cattle from Sokoto State, may be due to the fact that the sampling took place during the late dry season, while the adult population of this tick species tends to peak during the wet season. Accordingly, this tick was previously documented in cattle from Sokoto, surveyed between January and December [19] and February and July [21]. This species is indeed known to be widespread across Nigeria [8, 9, 16, 19, 21], being responsible for the transmission of several bovine TBPs such as *E. ruminantium*, *Dermatophilus congolensis, T. mutans* and *T. velifera* (reviewed in [8]). In this study, however, none of the three *A. variegatum* specimens screened was positive for Apicomplexan DNA.

One (female) specimen of *Rh. (Bo.) decoloratus* was also collected from a cow from Zamfara State. Known as the vector of *B. bigemina, A. marginale* and *A. centrale* [23], this species is considered the most prevalent boophilid tick infesting cattle in North-Central Nigeria [8, 13]. As for *A. variegatum*, its low abundance can be due to the seasonality of this tick, peaking during the wet season [8, 16], rather than when this survey took place.

*Rhipicephalus (Boophilus) microplus* was not recorded in any of the cattle sampled in this study, suggesting that the spread of this invasive tick, likely introduced in Nigeria through cattle from Benin [9] and so far identified in North-Eastern (i.e. Borno, Taraba and Yobe States) [9–11], North-Central (i.e. Kwara State) [9], South-Eastern (i.e. Enugu State) [12] and South-Western Nigeria (i.e. Ogun and Ondo States) [9], has probably not reached the North-West of the country. In Nigeria, indeed, this tick species can be expected to occur mostly in the southern regions, with higher relative humidity and annual precipitation compared to the drier northern States [9].

The preponderance of male rather than female specimens for most tick species collected in cattle from Zamfara State is an accordance with the existing literature (reviewed in [8]). Female ticks are indeed more frequently groomed off by cattle and tend to parasitise them for shorter periods compared to adult males [8]. The detection of more female than male specimens among the ticks collected in Sokoto State can be attributed to the context where the sampling took place. With sampling sites being rather crowded cattle markets, it is possible that our collection may have somehow privileged female ticks, that are indeed more conspicuous thus more easily identifiable with the naked eye, than male ones.

This study provides the first report of *T. annulata* in Nigeria. Tropical theileriosis is recognised as one of the most economically important diseases of livestock across North Africa and much of Asia [67], and its presence in SSA, where livestock productivity is already severely compromised by endemic parasites and pathogens [3], is an additional concern. In Africa, *T. annulata* has so far been detected in eight countries, mostly in the northern (i.e. Morocco, Algeria, Tunisia and Egypt) and eastern part of the continent (i.e. Sudan, South Sudan and Ethiopia) [67], with only Mauritania accounting for the locality records reported so far for West Africa [67–70].

Our detection of *T. annulata* DNA in ticks collected off 18 (of 35 tested) cattle in three different markets in Sokoto as well as off two (of seven tested) cattle in one village in Zamfara suggests it may be established in North-Western Nigeria. Considering that a significantly greater proportion of ticks and cattle from Sokoto contained *T. annulata* DNA than in Zamfara, it is likely that the parasite is more prevalent in the former State. That Sokoto is a likely port of introduction, be it recent or not, and circulation of this pathogen, is not unexpected. The State of Sokoto is indeed a major centre for livestock (including camels) trade in the region, attracting farmers and pastoralists not just from North-Western Nigeria, but also from neighbouring Niger and further afield in the Sahel and Saharan regions [71, 27]. The importance of trans-border trade as routes of entry of exotic ticks and TBPs into Nigeria has been established in the South-West of the country [72], with no information being available for the North of the country. Further characterisation of the *T. annulata* populations detected in Sokoto using, for example, previously described polymorphic markers [73] and comparison of these data with those obtained elsewhere in the parasite’s range, may help pinpoint their provenance.

It is currently unclear if *T. annulata* has been long-established in North-Western Nigeria or if it is a recent arrival. Indeed, that tropical theileriosis has not been reported before in Nigeria may reflect absence or a low awareness of it as a clinically apparent disease. The latter may also have resulted from endemic stability in the region coupled with low susceptibility of local cattle breeds to clinical disease, as reported elsewhere [74, 75]. Similarly, surveys carried out in cattle Mauritania, during the wet and the dry seasons, did not reveal any clinical case [69], with Zebus in the southern part of the country being found to have high specific antibody titres to *T. annulata* [68]. Nevertheless, a fatal case of bovine theileriosis was reported in a Friesian cow born in a dairy farm in Nouakchott, originally established with cattle imported from France, where *H. dromedarii* was the only tick species retrieved [68]. Surveys of local cattle are therefore urgently required to further exploration of the epidemiology of this infection and assess its clinical importance in cattle reared in North-Western Nigeria and potentially in other neighbouring regions. Importantly, the emergence and spreading of resistance to the most widely used theilericidal compound on the market (i.e. buparvaquone) in *T. annulata* strains circulating in Northern and Eastern Africa [67], further highlights the need for future epidemiological investigations.

Moreover, given that the susceptibility of camels to *T. annulata* has been established and infections appear common elsewhere [76], they too should be surveyed to explore what role they play in the natural cycle of *T. annulata* in the area. A small survey of TBPs in camels in Sokoto previously detected *Theileria* species, but not specifically *T. annulata* [77].

In the present study, five tick species were detected positive for *T. annulata* DNA, namely *H. dromedarii*, *H. impeltatum*, *H. rufipes*, *H. truncatum* and *H. impressum*. Of the four *Hyalomma* species implicated in the transmission of *T. annulata* in Africa (i.e. *H. scupense, H. anatolicum, H. dromedarii* and *H. lusitanicum*) [67], only *H. dromedarii* was encountered on cattle in this study, and at far greater abundance in Sokoto than Zamfara. Thus, it can be hypothesised that *H. domedarii* could be the most probable vector of *T. annulata* in North-Western Nigeria. However, other agro-ecological determinants, such as immature ticks’ host availability and husbandry practices may also underlie the inter-State differences we observed. Due to its capacity to withstand very hot and dry habitats, *H. dromedarii* is thought to have a strong comparative advantage over other endemic tick African species, in regions where climate change is expected to enhance the environmental aridity [67]. These characteristics, coupled with the vulnerability to climate change of North-Western Nigeria, may cause an increase of the prevalence of theileriosis in cattle from the region, in case *T. annulata* was only recently herein introduced and if the local vectorial competence of *H. dromedarii* was confirmed. Indeed, by creating new suitable habitats for *H. dromedarii*, the expansion of arid areas due to climate change, may lead the infection to spread across North-Western Nigeria, together with its presumable vector.

In Mauritania, *H. dromedarii* was proposed as the natural vector of *T. annulata* [68–70], although it was also noticed that the serological prevalence in cattle increased in areas with more diversified tick fauna, suggesting the potential implication of other species such as *Rhipicephalus evertsi evertsi* [68, 69]. This tick species was previously recorded in cattle in two surveys from Sokoto State (with a prevalence of 2.6% and 5.5%, respectively) [19, 21], as well as from south-eastern (i.e. Oyo State) [78] and eastern regions (i.e. Adamawa State) [79]. Therefore, the potential participation of *Rh. e. evertsi* in the epidemiology of *T. annulata* in North-Western Nigeria could not be excluded and should be further investigated. Future studies aiming to assess the occurrence of this tick species in Zamfara, are also advisable, especially considering that, besides Sokoto, the latter borders also with the States of Katsina and Kaduna, where *Rh e. evertsi* was recently identified in horses [80].

Like in this study, a few (n=2/30; 7%) partially fed *H. rufipes* ticks collected from cattle in the Gorgol region in Mauritania, were also found positive for *T. annulata* at PCR, although at a much lower prevalence compared to *H. dromedarii* (n=17/30; 57%) [70]. The vectorial competence of *H. rufipes* has indeed been demonstrated experimentally through studies on cattle, in which transstadial transmission (i.e. from nymph to adult) occurred [81].

Nevertheless, this tick is regarded as an unlikely vector of *T. annulata* in field conditions [81], since its immature stages usually parasitise ground-feeding birds [82]. Nymphal stages of *Hyalomma impeltatum* were also showed to transmit *T. annulata* in experimental conditions in Sudan [83]. Yet, this tick species is not considered a relevant vector of *T. annulata* either [83], because its immature stages mostly feed on rodents, hares and birds [84].

Although our study detected *T. annulata* DNA in several *Hyalomma* species, all but one specimen were partially fed or near replete, thus results cannot be interpreted as an indication of vector competence. Moreover, the not infrequent detection of *T. annulata* DNA in multiple ticks infesting the same individual cow underlines also the possible presence of this pathogen in the bovine blood. Interestingly, the only unfed tick specimen that was found positive in this survey was a male *H. dromedarii*, this being a further suggestion of the possible implication of this species in the transmission of *T. annulata* in the region. The state of repletion of male specimens of this tick is indeed identifiable through the examination of the position of the subanal plates (in fed individuals they protrude beyond the posterior margin of the body) [51].

## Conclusions

This study confirmed the presence of several tick species in North-Western Nigeria, most of which were previously documented in cattle and camels in Northern Nigeria and provides the first locality records for Zamfara State. The preponderance of *H. dromedarii* in cattle from Sokoto rather than Zamfara, coupled with the time of the year when the sampling took place (i.e. late dry season), highlights the suitability of this tick species for the arid environments of the Sahelian belt [51]. The potential expansion of drylands favoured by the ongoing climatic changes may potentially lead *H. dromedarii* to become more prevalent in the future in neighbouring southbound States (e.g. Zamfara, Kebbi and Kaduna). This warrants constant monitoring of the tick fauna infesting livestock of great economic relevance in Northern and Central Nigeria, such as ruminants and camels, including also cattle newly introduced from Niger or Chad for commercial purposes.

The occurrence of *H. turanicum*, recorded for the first time in Nigeria indicates a distribution of this tick beyond Northern Africa. Further studies aiming to better understand its occurrence and potential vectorial role, in SSA and Nigeria, would be advisable.

We also presented strong evidence for the presence of *T. annulata* in North-Western Nigeria and demonstrate its carriage by a range of primarily fed ticks collected off cattle. These observations pave the way for further epidemiological studies to clarify the transmission of *T. annulata* infections in the region and demand veterinary investigation of their impact on livestock well-being and productivity. Our findings and the existing literature suggest *H. dromedarii* as most probable vector in the area, although other species (e.g. *Rh. e. evertsi)* may also participate in the epidemiology of *T. annulata* infection.

Whether this infection has been newly introduced in Nigeria through trade cattle from Sahelian countries or whether it has long been endemically established in the local Zebu population, its presence represents a serious threat to the development of the livestock sector in North-Western Nigeria, given the severity of tropical theileriosis in crossbreed and exotic cattle species. For these reasons, this study’s findings need to be disseminated among veterinary authorities and livestock professionals in the country, to raise awareness on this tick-borne infection.

## Declarations

### Ethics approval and consent to participate

Approval for this study was obtained from the ethics committee of the University of Salford, Manchester, UK. At each study site in Anka (Zamfara), cattle owners (i.e. Fulani pastoralists) were approached and informed about the aims and methods of the study before being asked for informed consent. In northern Sokoto, the project was first discussed with Miyetti Allah, the local Fulani association that manages cattle markets in the region. Subsequently, cattle sellers in these markets were approached as described above and their informed consent requested.

### Authors’ contributions

AHM, VL, BMA, KJB and RJB conceived of the study and participated in its design. AHM, BMA, VL and RJB coordinated the field activities and tick collection. AHM and VL performed tick identification. AHM, BMA and RJB carried out the molecular analysis. VL and RJB took care of the statistical analysis. AHM, VL and RJB wrote the paper. All authors read and approved the final manuscript.

## Acknowledgements

We thank the Nigerian Petroleum Technology Development Fund for funding HAM’s MSc studies at the University of Salford, during which this research was performed. We also express appreciation to Adamu Samaila and his team for helping with the community sensitisation and sample collection. We are also grateful to all local authorities, cattle keepers and sellers involved in this study for their kind collaboration. Fabio Di Chio is also thanked for his kind assistance with preparation of the Figure.

## Authors’ information

AHM and VL contributed equally to this article.

